# Dose Escalation for Locally Advanced Pancreatic Cancer: How High Can We Go?

**DOI:** 10.1101/324129

**Authors:** Lauren E. Colbert, Neal Rebueno, Shalini Moningi, Sam Beddar, Gabriel Sawakuchi, Joseph Herman, Albert C. Koong, Prajnan Das, Emma Holliday, Eugene J. Koay, Cullen M. Taniguchi

## Introduction

Locally advanced pancreatic cancer (LAPC) is particularly difficult to treat given that it exhibits a poor response to chemotherapy and is by definition unresectable. A large proportion of patients experience significant morbidity and mortality due to local progression. As systemic therapy improves with the advent of gemcitabine/ abraxane^1^ and FOLFIRINOX^2^, the burden of local disease will only increase. Standard radiation therapy doses have failed to improve survival, which is not unexpected given anatomical and technical limitations. Although the LAP-07 trial^3^ failed to show an overall survival benefit to radiation at standard doses, it did show benefits in terms of local control and time off of chemotherapy. At higher radiation doses, however, our institutional data suggests improved overall survival (17.8 months vs 15 months)^4^ and recurrence-free survival (10.2 months vs 6.2 months), in addition to decreased acute grade 3+ gastrointestinal toxicity (1% vs 14%)^5^ even at higher doses (BED>70Gy) with advanced radiation delivery techniques including use of 4-dimensional computed tomography, breath-hold technique, and image-guided radiation therapy.

Given these initial retrospective data, we initiated a phase I/II adaptive dose escalation trial using image guided stereotactic body radiation therapy (SBRT) for LAPC at our institution (ClinicalTrials.gov Identifier: NCT03340974) to determine the clinical maximum tolerated dose using SBRT. In preparation for the activation of this protocol, we undertook a dosimetric feasibility study to determine a) whether all patients treated with IMRT could have also been planned with SBRT and b) the maximum feasible delivered biological effective dose (BED) using SBRT while maintaining standard organ at risk (OAR) constraints to gastrointestinal (GI) mucosa with daily imaging, motion management, and treatment planning techniques that would be available at most academic centers.

## Methods

### Patient Selection

The first ten sequential SBRT patients treated at our institution using SBRT at a dose of 40Gy in 5 fractions or 36Gy in 5 fractions (SD-SBRT) were selected for this study. Ten patients originally treated with dose-escalated hypofractionated IMRT (DE-IMRT; 67.5Gy in 15 fractions) were randomly selected from our previously published cohort in order to obtain a fair distribution of patients. All patients received 4 to 6 months of standard induction chemotherapy and on restaging exhibited unresectable disease based on a multidisciplinary review of computed tomography (CT) images using standard criteria^6^. Patients previously treated with DE-IMRT generally had tumors located >5mm from GI mucosa (OAR’s) with no predefined size limit, while SD-SBRT patients had tumors <4cm in maximal dimension with no evidence of duodenal invasion on imaging or endoscopy.

### Immobilization and Simulation

SD-SBRT patients had multiple fiducial markers implanted prior to simulation. DE-IMRT and DE-SBRT patients were immobilized using upper body vac-lock cradles. All patients were simulated NPO for three hours, using intravenous (IV) contrast and inspiration breath hold (IBH) with 5-6 scans for reproducibility. Typically two IBH CT scans were acquired without IV contrast followed by 3-4 IBH CT scans acquired after IV contrast injection performed in intervals of approximately 30 s between scans beginning 30 seconds after IV contrast administration. Simulation technique and example IBH CT scans are described in Figure 1.

**Figure 1.**
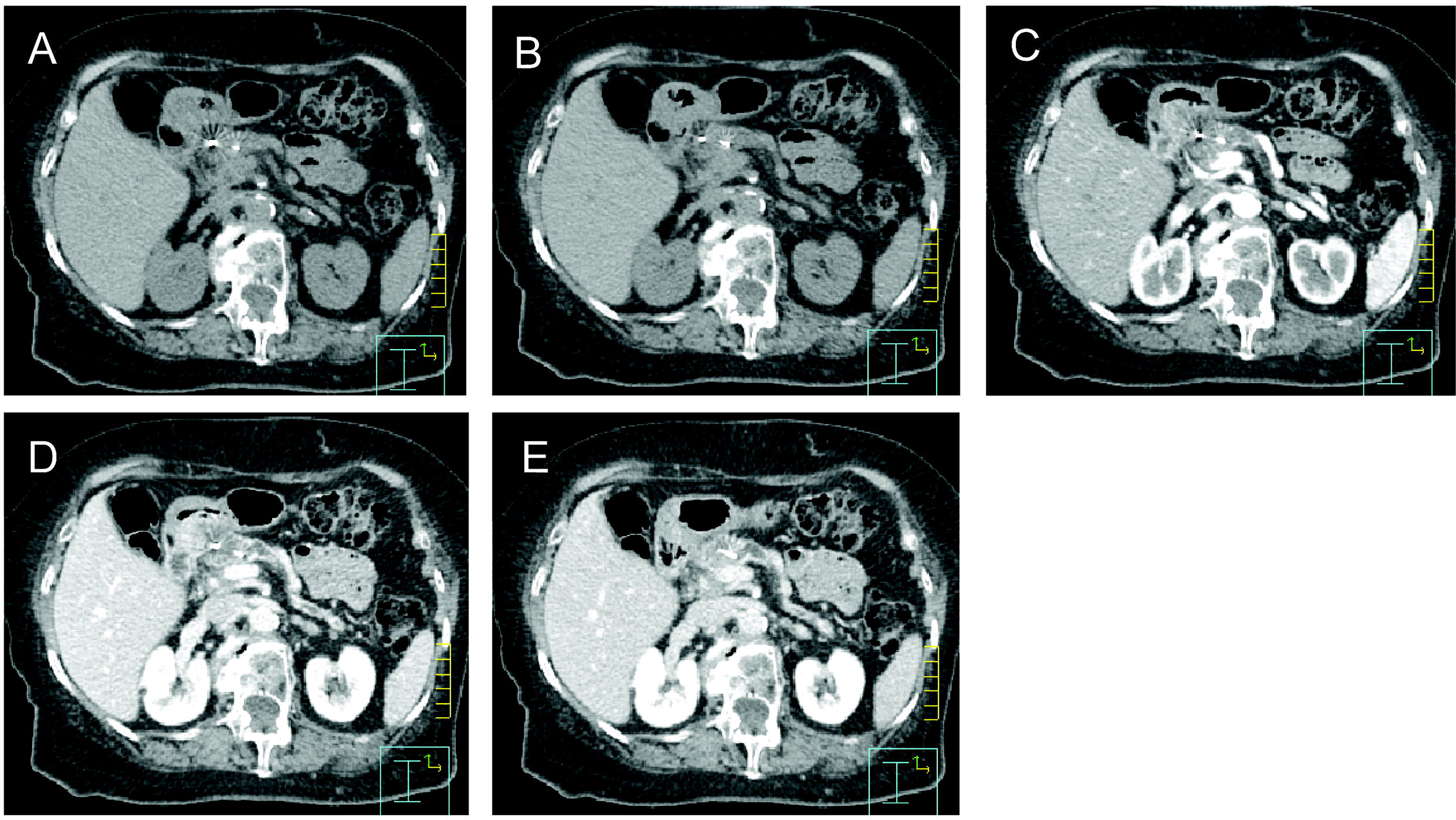
Serial BH images post contrast. Using a standard contrast enhanced protocol. Representative image from same axial slice in same patient at: A) 30s post-contrast infusion, B) 60s postcontrast infusion, C) 90s post-contrast infusion, D) 120s postcontrast infusion, and E) 150s post-contrast infusion

### Target Delineation and Creation of Simultaneous Integrated Boost (SIB)

All tumors from DE-IMRT and SD-SBRT patients were re-contoured and validated by two separate physicians (LE and CT) with identical targets using commercial treatment planning software (Phillips Pinnacle version 9.10). The IBH CT images were used to contour an integrated GTV (iGTV) and integrated OAR structures (iDuodenum, iStomach and iSmallBowel; Figure 2a). This is a similar concept to a respiratory internal target volume (ITV), created by accounting for physiologic movement of a target. The iOAR structures were uniformly expanded by 5mm expansion to create a GI mucosa planning risk volume (GI_PRV). Example contours are given in Figure 2.

**Figure 2.**
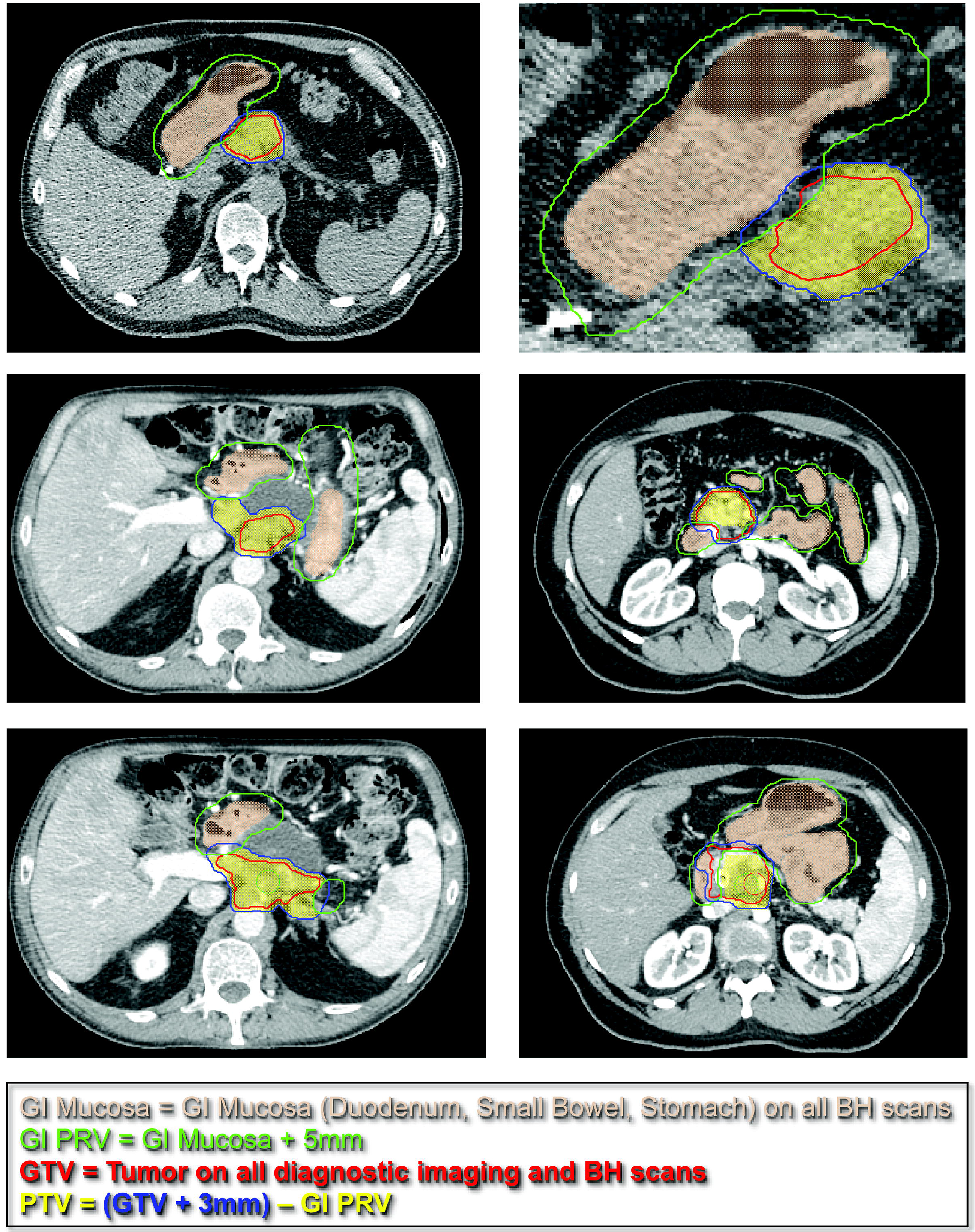
Target and OAR contouring technique using BH scans.

For each DE-IMRT and SD-SBRT, we attempted to created a dose-escalated SBRT (DE-SBRT) plan with an SIB to 40 Gy (8Gy/fx) and 70Gy (14Gy/fx) in 5 fractions. The PTV_40 was created by adding 3mm to the iGTV and subtracting the GI_PRV. A PTV_70 was created from the iGTV with 3mm contraction in 3 dimensions. DE-IMRT patients were re-planned for a 40Gy in 5 fractions SBRT plan and a 40Gy in 5 fraction plan with an increased simultaneous integrated boost (SIB) dose until the highest possible dose was reached (up to 70 Gy), while maintaining set OAR constraints (Table 1). The ten SD-SBRT patients were re-planned with a DE-IMRT plan and a DE-SBRT plan with an increased SIB. A tumor-vessel interface (TVI) has been included in recent trials to escalate dose to the SMA/ tumor interface. This was not used in our study to simplify plan comparisons, but areas of increased dose were pushed posteriorly to vessel interface rather than anteriorly whenever possible.

**Table 1.**
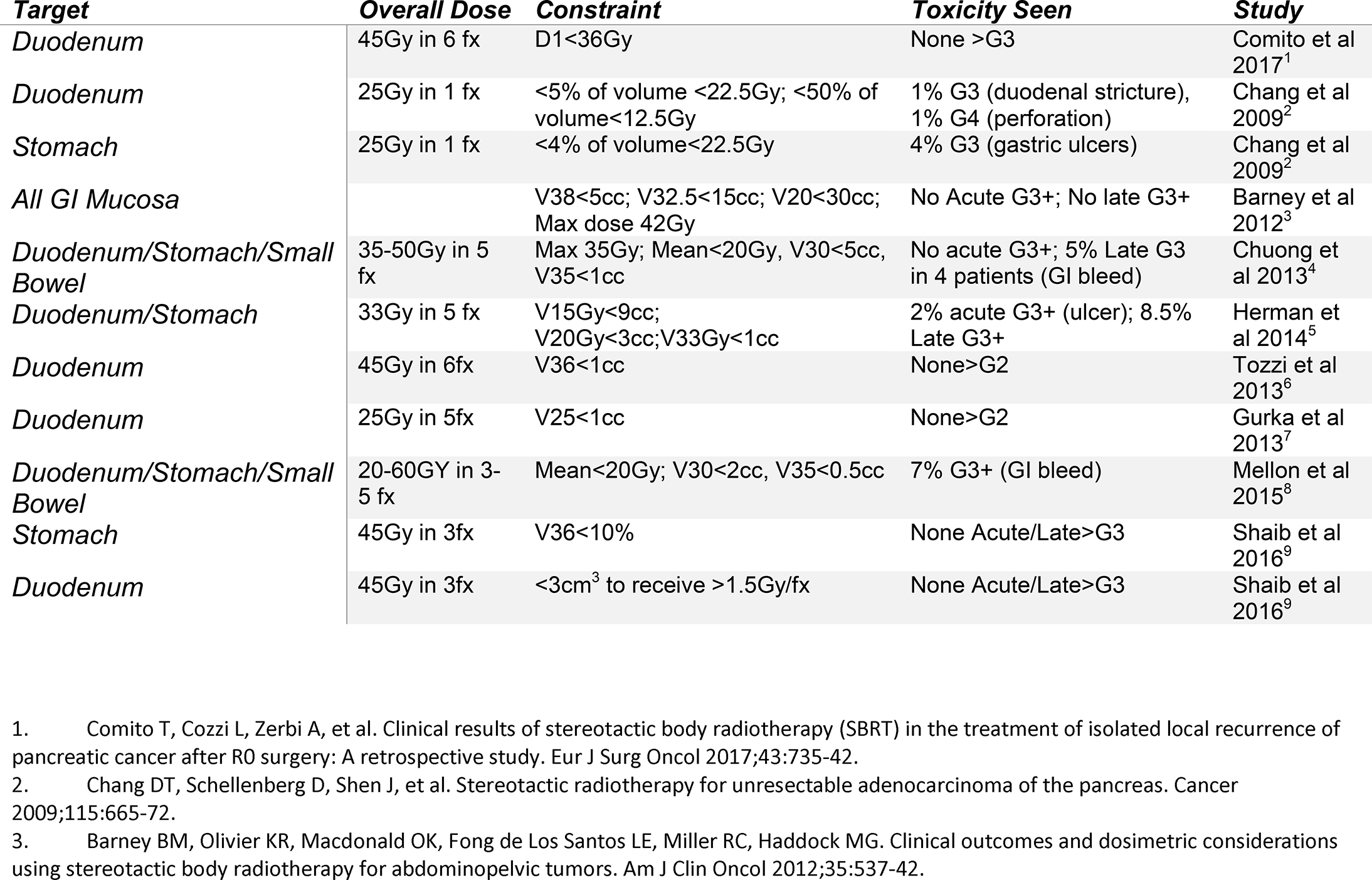
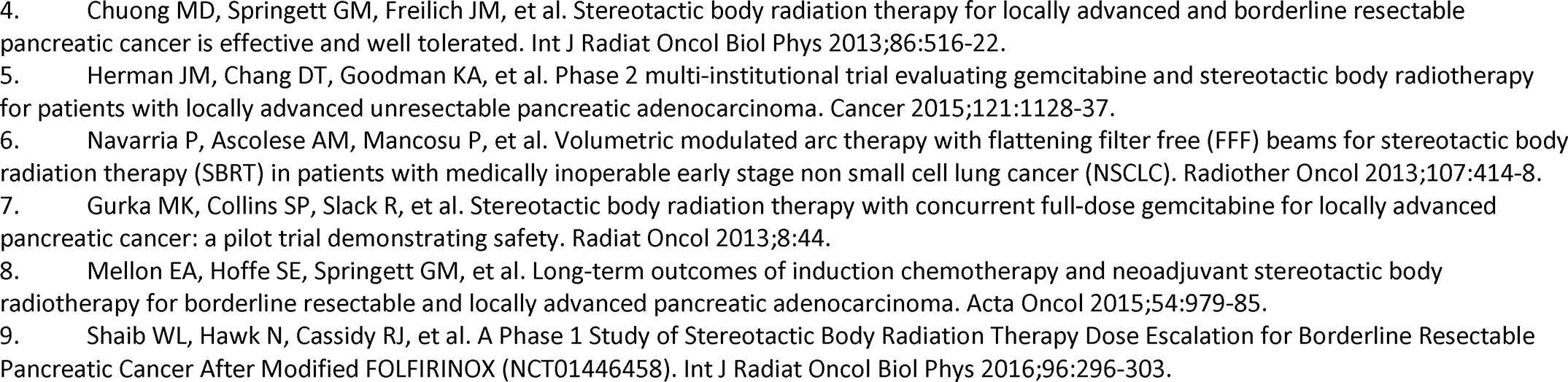
Previously Reported Dose Constraints and Associated Toxicities

| **Target** | **Overall Dose** | **Constraint** | **Toxicity Seen** | **Study** |
| --- | --- | --- | --- | --- |
| *Duodenum* | 45Gy in 6 fx | D1<36Gy | None >G3 | Comito et al 2017<sup>1</sup> |
| *Duodenum* | 25Gy in 1 fx | <5% of volume <22.5Gy; <50% of volume<12.5Gy | 1% G3 (duodenal stricture), 1% G4 (perforation) | Chang et al 2009<sup>2</sup> |
| *Stomach* | 25Gy in 1 fx | <4% of volume<22.5Gy | 4% G3 (gastric ulcers) | Chang et al 2009<sup>2</sup> |
| *All GI Mucosa* |  | V38<5cc; V32.5<15cc; V20<30cc; Max dose 42Gy | No Acute G3+; No late G3+ | Barney et al 2012<sup>3</sup> |
| *Duodenum/Stomach/Small Bowel* | 35-50Gy in 5 fx | Max 35Gy; Mean<20Gy, V30<5cc, V35<1cc | No acute G3+; 5% Late G3 in 4 patients (GI bleed) | Chuong et al 2013<sup>4</sup> |
| *Duodenum/Stomach* | 33Gy in 5 fx | V15Gy<9cc; V20Gy<3cc; V33Gy<1cc | 2% acute G3+ (ulcer); 8.5% Late G3+ | Herman et al 2014<sup>5</sup> |
| *Duodenum* | 45Gy in 6fx | V36<1cc | None>G2 | Tozzi et al 2013<sup>6</sup> |
| *Duodenum* | 25Gy in 5fx | V25<1cc | None>G2 | Gurka et al 2013<sup>7</sup> |
| *Duodenum/Stomach/Small Bowel* | 20-60GY in 3-5 fx | Mean<20Gy; V30<2cc, V35<0.5cc | 7% G3+ (GI bleed) | Mellon et al 2015<sup>8</sup> |
| *Stomach* | 45Gy in 3fx | V36<10% | None Acute/Late>G3 | Shaib et al 2016<sup>9</sup> |
| *Duodenum* | 45Gy in 3fx | <3cm<sup>3</sup> to receive >1.5Gy/fx | None Acute/Late>G3 | Shaib et al 2016<sup>9</sup> |

1. Comito T, Cozzi L, Zerbi A, et al. Clinical results of stereotactic body radiotherapy (SBRT) in the treatment of isolated local recurrence of pancreatic cancer after R0 surgery: A retrospective study. *Eur J Surg Oncol* 2017;43:735-42. 2. Chang DT, Schellenberg D, Shen J, et al. Stereotactic radiotherapy for unresectable adenocarcinoma of the pancreas. *Cancer* 2009;115:665-72. 3. Barney BM, Olivier KR, Macdonald OK, Fong de Los Santos LE, Miller RC, Haddock MG. Clinical outcomes and dosimetric considerations using stereotactic body radiotherapy for abdominopelvic tumors. *Am J Clin Oncol* 2012;35:537-42.
4. Chuong MD, Springett GM, Freilich JM, et al. Stereotactic body radiation therapy for locally advanced and borderline resectable pancreatic cancer is effective and well tolerated. *Int J Radiat Oncol Biol Phys* 2013;86:516-22. 5. Herman JM, Chang DT, Goodman KA, et al. Phase 2 multi-institutional trial evaluating gemcitabine and stereotactic body radiotherapy for patients with locally advanced unresectable pancreatic adenocarcinoma. *Cancer* 2015;121:1128-37. 6. Navarria P, Ascolese AM, Mancosu P, et al. Volumetric modulated arc therapy with flattening filter free (FFF) beams for stereotactic body radiation therapy (SBRT) in patients with medically inoperable early stage non small cell lung cancer (NSCLC). *Radiother Oncol* 2013;107:414-8. 7. Gurka MK, Collins SP, Slack R, et al. Stereotactic body radiation therapy with concurrent full-dose gemcitabine for locally advanced pancreatic cancer: a pilot trial demonstrating safety. *Radiat Oncol* 2013;8:44. 8. Mellon EA, Hoffe SE, Springett GM, et al. Long-term outcomes of induction chemotherapy and neoadjuvant stereotactic body radiotherapy for borderline resectable and locally advanced pancreatic adenocarcinoma. *Acta Oncol* 2015;54:979-85. 9. Shaib WL, Hawk N, Cassidy RJ, et al. A Phase 1 Study of Stereotactic Body Radiation Therapy Dose Escalation for Borderline Resectable Pancreatic Cancer After Modified FOLFIRINOX (NCT01446458). *Int J Radiat Oncol Biol Phys* 2016;96:296-303.

### Treatment Planning and Evaluatio

Planning was performed in one of the IV contrast IBH CT scans without density override for the IV contrast. 40Gy was prescribed to the PTV_40 with a SIB technique prescribing 70Gy to the PTV_70. There was no minimum coverage requirement for the iGTV, but >95% coverage was requested for the PTVs. OAR’s were prioritized during IMRT planning over target coverage. Successful plans typically had between 7 and 12 coplanar beam angles. VMAT was acceptable if available for treatment delivery with breathhold. Example treatment plans are presented in Figure 3. Based on toxicities observed in previous SBRT trials and the associated constraints (Table 1), dose constraints defined for planning purposes (Table 2) were: Duodenum V20<20cc, V35<1cc, Dmax<40Gy; Small Bowel V20<20cc, V35<1cc, Dmax<40Gy; Stomach V20<20cc, V35<1cc, Dmax<40Gy.

**Figure 3.**
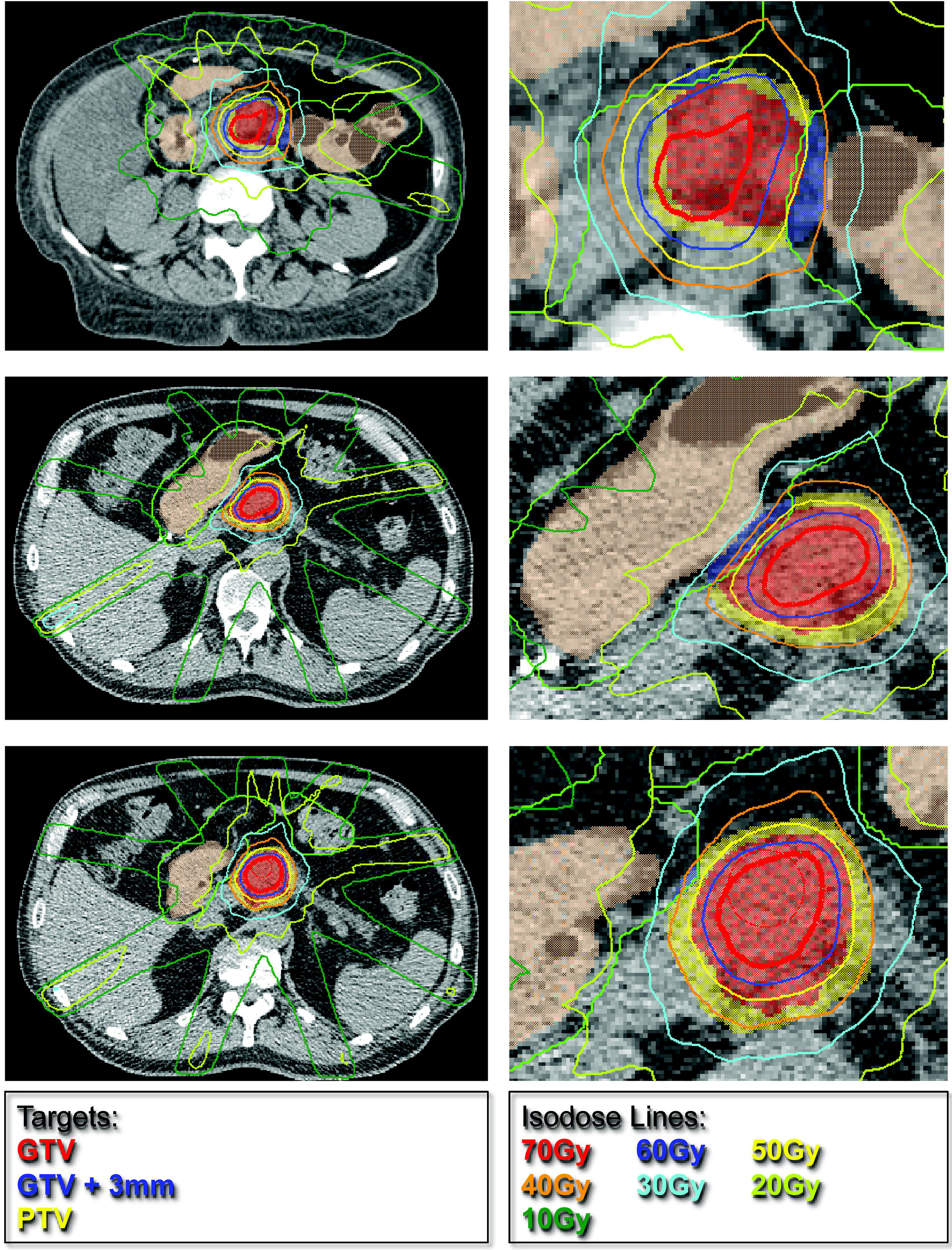
Sample plans for deSBRT patients. Each row is a different set of patients. Left column is full axial slice and accompanying right column is zoomed in on areas of interest.

**Table 2.** Pre-study and post-study Dose Constraints for Dose Escalated SBRT

| ***Target*** | ***Constraint Used for Planning*** | ***Adapted Constraint*** |
| --- | --- | --- |
| *Duodenum* | V20<20cc | V20<30cc |
|  | V35<1cc* | V30<3cc |
|  | Dmax<40Gy | V35<1cc |
|  |  | V40<0.5cc |
| *Small Bowel* | V20<20cc | V20<15cc |
|  | V35<1cc* | V30<1cc |
|  | Dmax<40Gy | V35<0.1cc |
|  |  | V40<0.5cc |
| *Stomach* | V20<20cc | V20<20cc |
|  | V35<1cc* | V30<2cc |
|  | Dmax<40Gy* | V35<1cc |
|  |  | V40<0.5cc |
| *Kidneys* | V12<25%* | V12<25% |
| *Liver* | V12<50%* | V12<50% |
| *Spinal Cord* | V20<1cc* | V20<1cc |

\* Mandatory constraints

Conformality index was calculated using prescription isodose volume (PIV; 40Gy)/ prescription target volume (PTV) and homogeneity index was calculated using both D95/D5 and Dmax/Dmin formulas. Gradient index was calculated using 50% PIV / 100% PIV.

### Statistical Analysis

Descriptive statistics were generated for all targets and all OAR. The 90^th^ percentile was used as a minimum threshold for reasonably achievable OAR contstraints. A minimum coverage percent of 60 percent was defined as acceptable for a dose escalation level.

### Treatment Delivery

All DE-IMRT were treated in a Clinac 2100 EX (Varian Medical Systems, Palo Alto, CA) equipped with a CT on rails (CTOR) (Smart Gantry, GE Healthcare, Chicago, IL). SD-SBRT patients were treated in a Truebeam (Varian Medical System) or the Clinac 2100 EX. For CTOR treatments, patients were initially set up under IBH using radiopaque skin marks placed on the planning final isocenter followed by CTOR under IBH. In-house software (Court and Dong 2003, Med Phys 30:2750) was used to register the CTOR to the planning CT using a contour of the vertebral body located just posterior to the iGTV. The final targets were then manually aligned using the implanted fiducial markers. The auto registration software then provides the couch shifts relative to the radiopaque skin marks to align the patient to the treatment final isocenter. Finally, a MV image pair is acquired to verify the bony alignment and treatment is delivered. For the Truebeam treatments, patients were initially set up to the planning final isocenter then aligned to fiducials using a kilovoltage image pair with cone beam CT (CBCT) for verification. Schema for standard treatment delivery using CTOR is given in Figure 5.

## Results

Clinically acceptable DE-SBRT plans based on iGTV and PTV coverage and OAR constraints were generated for 100% of patients originally treated with DE-IMRT. Acceptable DE-IMRT plans were also generated for 100% of patients originally planned with SD-SBRT. For the maximum dose escalated SBRT plans (Figure 4), mean PTV volume was 58.8cc (range 8.7-171.91; ±41.3cc). Mean iGTV volume was 40.4cc (range 3.3-119.06; ±31.9). Mean conformality index was 0.98 (±0.24). Mean homogeneity index using Dmax/Dmin was 2.26 (±0.21) and using D95/D5 was 1.77 (±0.05). Mean PTV coverage by the 40Gy line was 97.6% (±0.02%). Mean iGTV coverage by 50Gy was 91% (±0.07%), by 60Gy was 61.3% (±0.08%) and by 70Gy was 24.4% (±0.05%). Maximum PTV coverage by 70Gy was 33%. Maximum PTV coverage by 60Gy was 77.5%. Distributions for conformality and homogeneity and gradient indices and target coverage are given in Figure 3. Distributions for V20, V30 and V35 for all OAR’s are given in Figure 3. The following OAR constraints were achieved for ≥90% of generated plans: Duodenum V20<30cc, V30<3cc, V35<1cc; Small Bowel V20<15cc, V30<1cc, V35<0.1cc; Stomach V20<20cc, V30<2cc, V35<1cc. V40<0.5cc was achieved for all OAR. These post-study dose constraints are listed in table 2.

**Figure 4.**
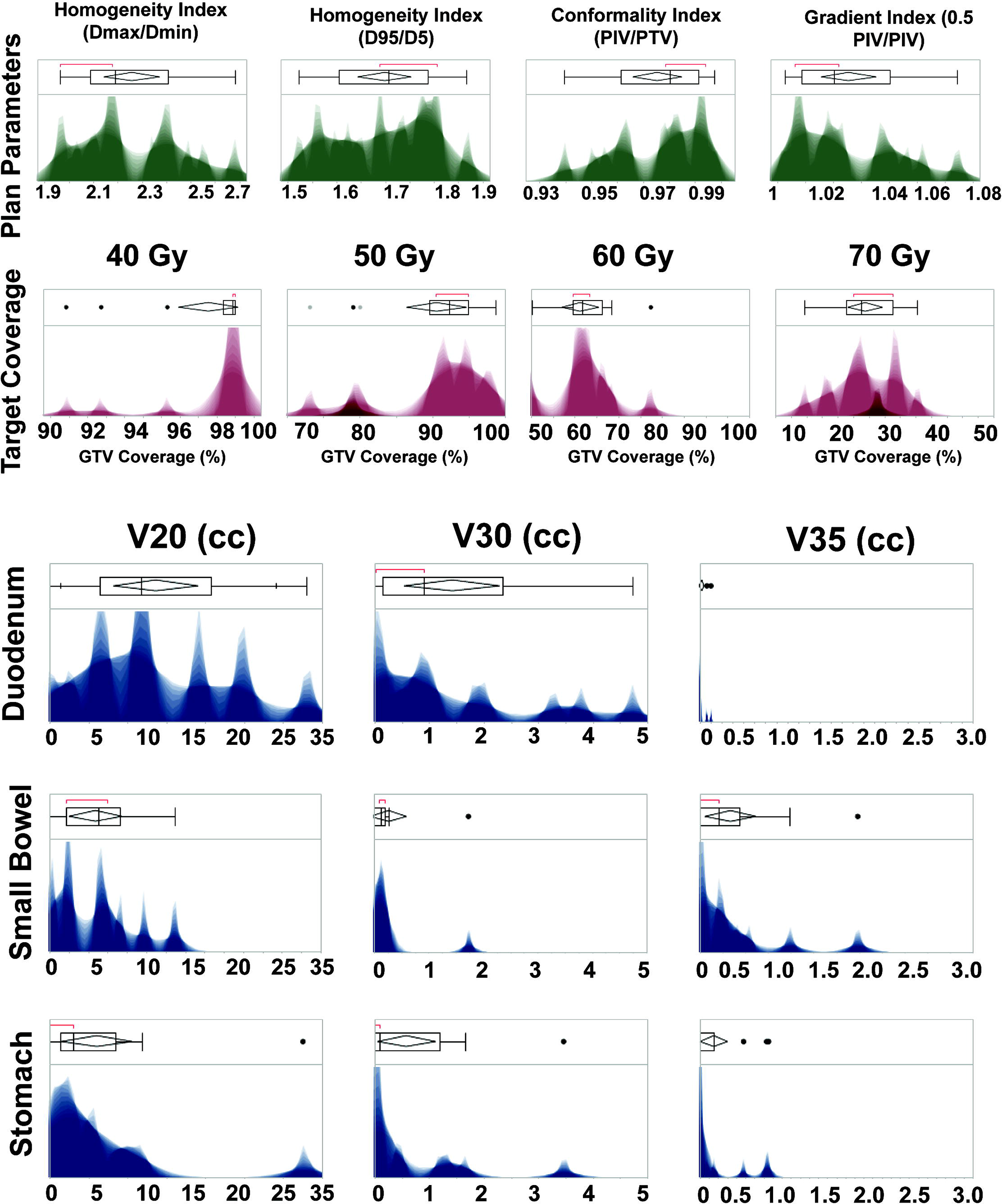
Distribution of OAR constraints and GTV/PTV coverage tradeoff.

**Figure 5.**
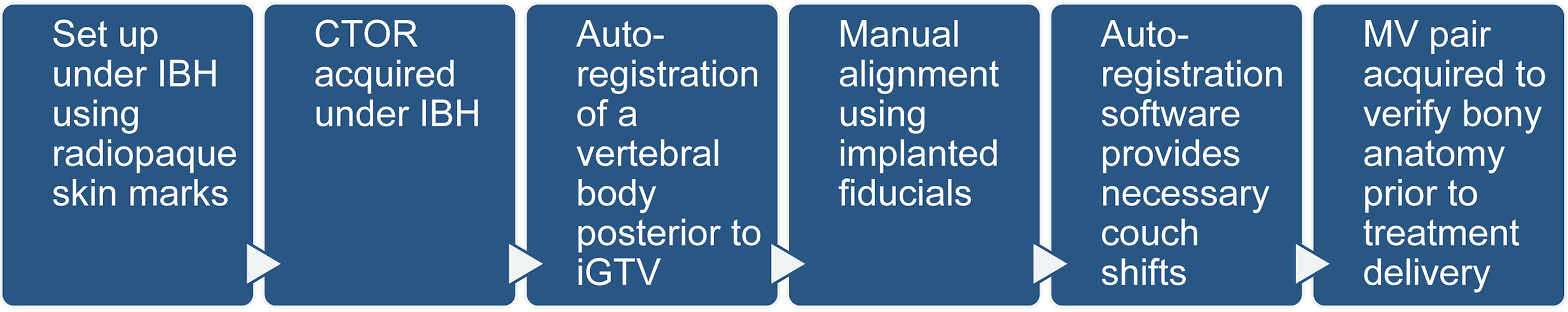
Typical Daily IGRT Process for DE-SRBT.

## Discussion

Dosimetrically, all patients treated with IMRT could also have been treated with SBRT, with similar (or improved) BED delivery. Given this, it is worth investigating SBRT techniques for patient convenience and continuity of systemic therapy. Based on the 90^th^ percentile used in the above data, dose escalation with an SIB technique to 60Gy in 5 fractions is achievable while maintaining acceptable target coverage and standard OAR constraints. We note that our study was enriched in patients with more favorable anatomy (uncinate and body tumors), but we believe that our approach and dose constraints would also apply to any patient that is eligible for pancreatic SBRT.

Our SIB approach covered the GTV between 60% and 80% by the highest doses while still maintaining >98% PTV coverage by the 40Gy line. Pancreatic tumors are particularly hypoxic at their core^7^, and delivering high doses to this hypoxic core may have a radiobiologic advantage despite not achieving full target coverage. This concept is similar to acceptance of dose heterogeneity within the GTV in other forms of SBRT^8^. Past SBRT trials, even with three and one fraction regimens have shown that duodenum, small bowel and stomach constraints of V20<30cc, V35<1cc, and max dose<40Gy are safe and well tolerated (Table 1). Using the above planning technique, these same dose constraints are achievable in 90% of cases, which supports the idea that dose escalation for LAPC is feasible and should be investigated in clinical trials. Our data provide a roadmap for other clinicians looking to achieve dose escalation up to 60Gy in 5 fractions for pancreatic cancer in the appropriate setting.

## Notes

**Conflicts of Interest:** None

